# Application of high-resolution landmark-free morphometrics to a mouse model of Down Syndrome reveals a tightly localised cranial phenotype

**DOI:** 10.1101/711259

**Authors:** Nicolas Toussaint, Yushi Redhead, Wei Liu, Elizabeth M. C. Fisher, Benedikt Hallgrimsson, Victor L.J. Tybulewicz, Julia A. Schnabel, Jeremy B.A. Green

## Abstract

Characterising phenotypes often requires quantification of anatomical shapes. Quantitative shape comparison (morphometrics) traditionally uses anatomical landmarks and is therefore limited by the number of landmarks and operator accuracy when landmarks are located manually. Here we apply a landmark-free method to characterise the craniofacial skeletal phenotype of the Dp1Tyb mouse model of Down syndrome (DS), validating it against a landmark-based approach. We identify cranial dysmorphologies in Dp1Tyb mice, especially smaller size and brachycephaly (front-back shortening) homologous to the human phenotype. The landmark-free phenotyping was less labour-intensive and required less user training than the landmark-based method. It also enabled mapping of local differences as planar expansion or shrinkage. This higher resolution and local mapping pinpointed reductions in interior mid-snout structures and occipital bones in this DS model that were not as apparent using a traditional landmark-based method. This approach could make morphometrics widely-accessible beyond traditional niches in zoology and palaeontology, especially in characterising mutant phenotypes.

## Introduction

Morphometrics, the quantitative comparison of biological shapes, is well established in the fields of palaeontology and evolutionary biology to quantify and understand morphological phenotypes (Cooke and Terhune, 2015). Landmark positions are recorded on digital two- or three-dimensional images (obtained by photography, X-ray or MRI methods) and their spatial distributions are then analysed through Euclidean distance matrix analysis (EDMA) or Procrustes Superimposition (PS) (Webster and Sheets, 2010). Morphometrics is less used in other fields, such as genetics and developmental biology. This may be because current morphometric methodologies, while powerful, have limitations. First, the number of landmarks always reflects a compromise between precision, which needs many anatomical landmarks to be located, and ease-of-use, which limits those numbers. Typically, some tens of landmarks are located manually, which takes anatomical knowledge and training and time. Second, an anatomical landmark may be absent from an individual due to natural variation, engineered mutation or pathology. Third, landmarks can be sparse in anatomical structures where they are hard to define: smooth surfaces do not have easily defined landmarks. Sparseness is a particular problem in soft tissues and embryos, with numerous featureless, curved surfaces. Semi-landmarks interpolated between landmarks (Andresen et al., 2000; Bookstein, 1997; Frangi et al., 2003) reduce this problem but still leave gaps (Palci and Lee, 2019). Fourth, manual landmark-based methods are inevitably susceptible to both inter- and intra-operator variability, which can be as big as the biological variability between subjects (Percival et al., 2014; Shearer et al., 2017; von Cramon-Taubadel et al., 2007). Together, these limitations suggest that there is a need for automated, ideally landmark-free, high-resolution methods. Landmark-free methods have been developed by the neuroimaging community to quantify the size and shape of the brain precisely because its relatively smooth shape hampers the definition of reliable landmarks (Bron et al., 2015; Routier et al., 2014) but these methods have yet to be applied more widely and have not been directly compared to the landmark-based approach.

One of the most common human dysmorphologies is the craniofacial phenotype associated with Down syndrome (DS). Individuals with DS, currently ∼1 in 800 births (Antonarakis, 2017), have characteristic features – flattened midface with low nose bridge, front-to-back shortened skull (brachycephaly) and slightly hooded eyelids (Korenberg et al., 1994). Although the craniofacial features affect everyone with DS, this phenotype is not well understood either genetically or developmentally. DS is caused by trisomy of human chromosome 21 (Hsa21) which carries 232 protein-coding genes (Ensembl genome assembly GRCh38) (Antonarakis, 2017; Lejeune et al., 1959). It is thought that the presence of a third copy of one or more of these genes (rather than just the higher chromosomal load) gives rise to the individual defects observed in DS, but the critical dosage-sensitive genes are not known (Lana-Elola et al., 2016; Lana-Elola et al., 2011; Watson-Scales et al., 2018).

To model DS, mouse strains have been engineered that carry an extra copy of each of the three regions of the mouse genome orthologous to Hsa21. These recapitulate at least some aspects of DS (Herault et al., 2017; Lana-Elola et al., 2016; Yu et al., 2010). Morphometrics applied to Ts65Dn (Hill et al., 2007; Richtsmeier et al., 2000; Richtsmeier et al., 2002) and Dp(16)1Yey mice (Starbuck et al., 2014) showed that trisomy of part or all of the Hsa21-orthologous region of mouse chromosome 16 (Mmu16) resulted in craniofacial dysmorphology which resembled the DS phenotype. The cranial dysmorphology in Dp1Yey mice was highly statistically significant (with multiple linear distances between landmarks differing statistically significantly from wild type in all regions measured) yet quantitatively subtle, with an average landmark-to-landmark distance difference of only 7% between mutant and wild-type (WT) control mice (Starbuck et al., 2014).

In this paper, we describe a convenient pipeline we have developed for landmark-free morphometric analysis based on an approach used for brain imaging (Durrleman et al., 2014). We compare our method to the traditional landmark-based morphometric approach, focusing on the characterisation of the craniofacial phenotype of the Dp1Tyb mouse model of DS which has an additional copy of the entire Hsa21-orthologous region of Mmu16 (Lana-Elola et al., 2016). We find that the landmark-free analysis gives separation by shape between Dp1Tyb and WT mice that is at least as clear as that achieved by landmark-based analysis, while delivering a number of operational advantages. We demonstrate a new tool (“local stretch” mapping) that avoids the need to separate scale changes from shape changes, and localises abnormalities in the DS model to cranial vault expansion and mid-face and occipital contraction.

## Results

### Landmark-based and landmark-free analysis of Dp1Tyb skulls

To phenotype the Dp1Tyb DS model skulls, we used micro-computed tomography (μCT) to acquire images of the skulls of 16-week old WT and Dp1Tyb mice. We carried out landmark-based analysis in the conventional way (Kristensen et al., 2008), marking the location of 68 landmarks on the cranium and 17 on the mandible (Supplementary Fig. 1). Crania and mandibles were analysed separately since their relative position varied from subject to subject. Landmarks for all crania and mandibles were aligned using Procrustes Superimposition, and these data were used for further statistical analysis of size and shape.

For the landmark-free approach we developed a pipeline based on previous approaches in morphometrics and neuroimaging (Fig. 1, Table 1, Supplementary Appendix 1). In brief, following thresholding to extract the skull structures from the μCT images, cartilaginous structures were removed (Supplementary Fig. 2) and the images segmented using bone density to separate the mandibles from the crania (step 1). Triangulated meshes were generated from the surfaces (including internal surfaces) of the cranium and mandible for all subjects (step 2), aligned (and scaled where appropriate – see below) (step 3). The meshes were used for the construction of an atlas (mean shape) for the crania and mandibles of the WT and Dp1Tyb skulls (step 4). Atlas construction was based on the Deformetrica algorithm (Durrleman et al., 2014) which works by defining a flow field (tensor) that conforms to its shape and quantifies deformations from it to each subject recorded as momentum vectors (momenta - see Methods and Supplementary Appendix 1). The initial output from this atlas consisted of the average mesh for the whole population (based on averaging the tensors), a set of control points corresponding to areas with the greatest variability between subjects, and momenta for each control point describing the directional variation of the shape from the average. The average mesh, the control points and the momenta were used for further statistical analysis, with the momenta applied to deform the population average mesh to generate average meshes for each of WT and Dp1Tyb groups preserving one-to-one correspondence of mesh vertices. We performed principal component analysis and used a multiple permutations test on a stratified k-fold cross validation classifier to test for significance (step 5). To control for overfitting (a risk when the number of measurements substantially exceeds the number of subjects), we compared the PCA difference vector magnitude between the two genotype groups with that of 1000 randomly scrambled groups. We found that the distribution was Normal and that the genotype difference vector was more than 3.5 standard deviations away from the mean vector of the 1000 scrambled groups for both cranium and mandible, thus showing that overfitting is unlikely to be a significant factor (data not shown).

**Table 1.** Summary of steps in the landmark-free analysis pipeline and chosen parameters at each step.

| Step | Sub-step | Function | Parameters | Tool | Reference |
| --- | --- | --- | --- | --- | --- |
| **Step 1.**<br>**Pre-processing** | Binary segmentation | Binary threshold | threshold = 10000 | FSL | (Jenkinson et al., 2012) |
|  |  | Closing | kernel size = 5 |  |  |
|  |  | Opening | kernel size = 3 |  |  |
|  |  | Clustering and extraction of largest component |  |  |  |
|  |  | Fill holes |  |  |  |
|  | Object parcelation | Watershed | watershedlevel = 0.22 | SimpleITK | (Yaniv et al., 2018) |
| **Step 2.**<br>**Mesh extraction** | Mesh extraction | Marching cubes |  | VTK | (Schroeder et al., 1998) |
|  |  | Smoothing | number of iterations = 20<br>relaxation factor = 0.6 |  |  |
| **Step 3.**<br>**Mesh Alignment** | Alignment points | Reference points used to align meshes |  | MITK | (Nolden et al., 2013) |
|  | Mesh Alignment | Centroid size |  | VTK/Numpy |  |
|  |  | vtkProcrustesAlignmentFilter | Rigid body or similarity | VTK |  |
| **Step 4.**<br>**Atlas** | Atlas construction | Atlas construction | data-sigma = 10<br>kernel width (template) = 2.5mm<br>kernel width (subject) = 2mm<br>number of time points = 10<br>max iterations = 150<br>number of threads = 19 | Deformetrica | (Gori et al., 2017) |
| **Step 5.**<br>**Statistical analysis** | Principal component analysis | Kernel PCA | number of components = 5 | Scikit-learn | (Pedregosa et al., 2011) |
|  |  | Kernel PCA lambdas (eigen values) |  |  |  |
|  |  | Support vector machine |  |  |  |
|  |  | Permutation test |  |  |  |
|  |  | Cross validation |  |  |  |
|  | Momenta projection / morph | Shooting | template kernel width = 2.0<br>deformation kernel width = 1.5 | Deformetrica |  |

**Figure 1.**
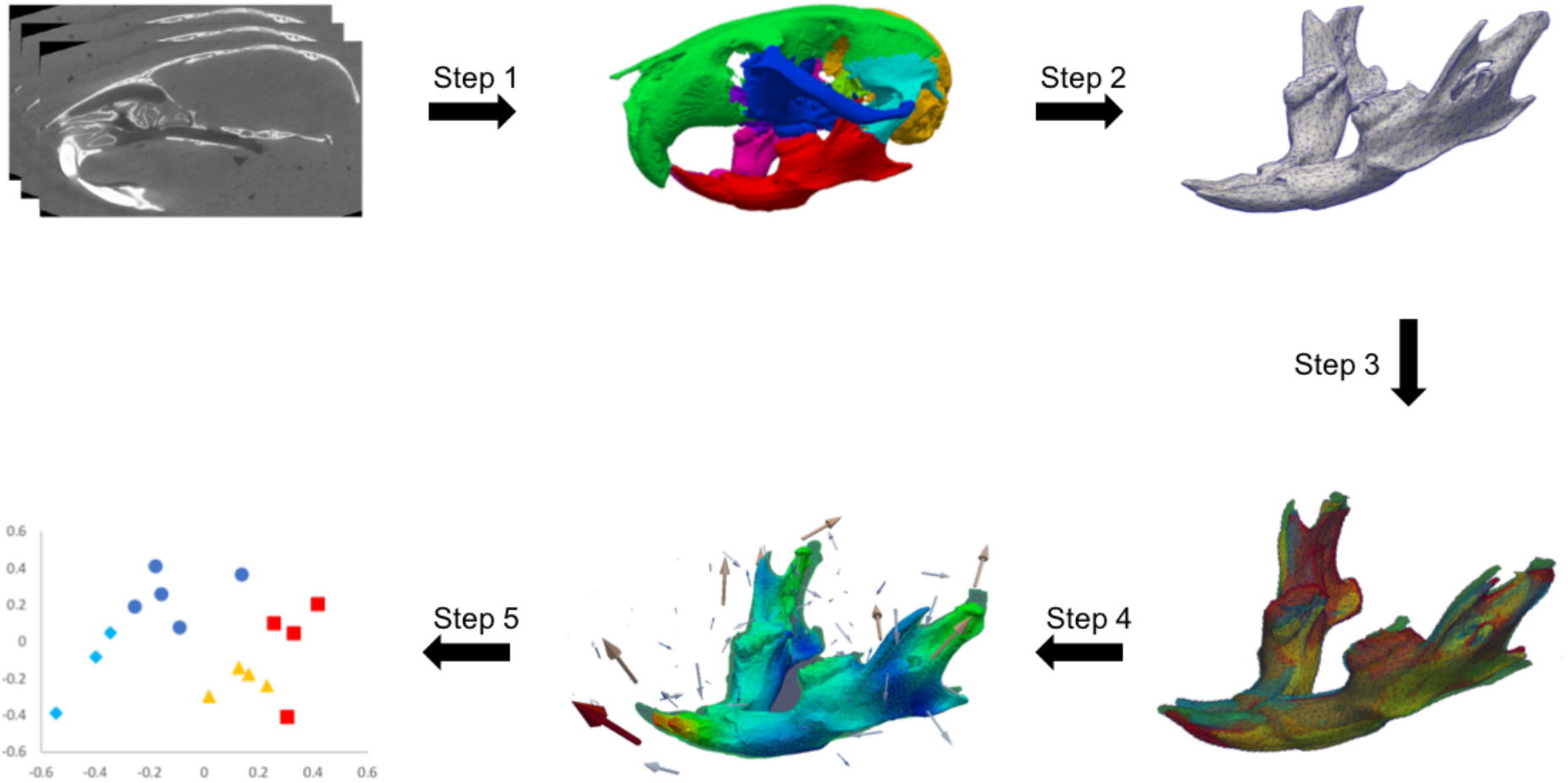
Stages of processing in the landmark-free approach. Step 1, extraction of region of interest. Initial thresholding of μCT image was used to make a binary mask and regions of the mask were separated by bone density using secondary thresholding, with some manual clean-up based on known anatomy. A region of interest (in this example the mandible in red) was chosen for further analysis. Step 2, mesh generated and decimated by a factor of 0.0125 to reduce data file size. Step 3, meshes of all subjects aligned either using rigid body alignment with no scaling or similarity alignment with scaling. Step 4, atlas construction. Step 5, statistical analysis and visualisation of shape data.

### Size differences: Dp1Tyb mice have significantly smaller crania and mandibles

We used centroid size (the mean absolute landmark distance from the landmark-defined centroid) to compare overall sizes of Dp1Tyb and sibling control specimens (Klingenberg, 2016). Landmark-based centroid size comparison showed that the crania and mandibles of Dp1Tyb mice were both significantly smaller than those of WT mice (Fig. 2A, C), recapitulating the overall reduction in skull size found in humans with DS and as well as in other models of DS (Hill et al., 2007; Richtsmeier et al., 2000; Richtsmeier et al., 2002; Starbuck et al., 2014; Suri et al., 2010). The landmark-free analysis also showed that Dp1Tyb crania and mandibles were significantly smaller (Fig. 2B, D). Both methods showed an approximately 7% reduction in centroid size of both cranium and mandible in Dp1Tyb mice (Supplementary Table). The landmark-free and landmark-based centroid sizes were different in absolute magnitude, unsurprisingly given the many extra measurements use in the landmark-free method (∼19,000 mesh vertices versus 68 landmarks for cranium and ∼16,000 vertices versus 17 landmarks for mandible).

**Figure 2.**
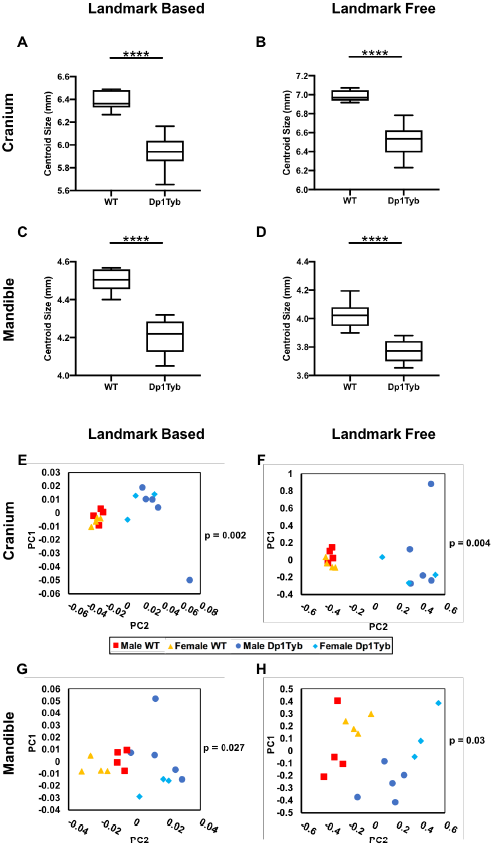
Decreased size and altered shape of Dp1Tyb crania and mandibles. **A-D** Centroid sizes of WT and Dp1Tyb crania (**A, B**) and mandibles (**C, D**) determined using landmark-based (**A, C**) and landmark-free (**B, D**) methods. Data shown as box and whiskers plots indicating the 25% and 75% centiles (box), range of all data points (whiskers) and the median (black line). Statistical significance was calculated using a two-tailed unpaired t-test; **** *P* < 0.0001. **E-H** Principal component analysis (first two components) of Procrustes aligned shapes of WT and Dp1Tyb crania (**E, F**) and mandibles (**G, H**) determined using landmark-based (**E, G**) and landmark-free (**F, H**) methods. Statistical significance of the differences between the WT and Dp1pTyb groups of samples was calculated using a permutation test, and *P*-values are shown on the plots. Sample size: n = 8 of each genotype.

### Shape differences: Dp1Tyb mice have altered crania and mandibles

Both the size difference and gross shape differences were clearly visualised by animated morphing between the mean shapes of WT and Dp1Tyb specimens (generated in the landmark-free pipeline) for the cranium and the mandible (Supplementary Videos 1, 2). The overall decrease in size going from WT to Dp1Tyb crania or mandibles was readily apparent and some shape changes could also be seen, although the latter were more subtle.

To quantify shape differences statistically, shape was separated from size by scaling the data to equalise centroid sizes (Procrustes alignment). To analyse residual shape differences between genotypes, we used principal component analysis (PCA). Both landmark-based and landmark-free methods showed a statistically significant differences in shape between Dp1Tyb and WT mice in both crania and mandibles (Fig. 2E-H). Plots of the first two Principal Components identified by the two different methods looked similar, with tighter clustering of specimens for cranium than for mandible.

### Shape difference localisation: Dp1Tyb mice recapitulate aspects of human DS craniofacial dysmorphology

To characterise the shape differences anatomically, we first, simply overlaid the mean landmark configurations from Dp1Tyb and WT crania and mandibles (Fig. 3A-F). Second, we applied an established thin-plate spline interpolation and comparison package (Morpho R - see Methods) to the landmark data to generate displacement heatmaps (Fig. 3G-L). Direct inspection revealed that the morphological differences between Dp1Tyb and WT skulls were broadly distributed and relatively subtle, consistent both with previously reported mouse DS models and with the human phenotype (Fischer-Brandies, 1988; Fischer-Brandies et al., 1986; Suri et al., 2010). The maps revealed that Dp1Tyb mice have a more domed neurocranium (cyan points at the top-right of Fig. 3A, dark red regions in Fig. 3G, H, J). The cranial doming in combination with the overall smaller size compared to WT mice constitute a net anteroposterior shortening in Dp1Tyb mice, i.e. brachycephaly, a predominant feature of the human DS phenotype. This method also indicated an almost unchanged cranial base (Fig. 3I) and some contraction (anterior movement) concentrated around the magnum foramen in the occipital bone of Dp1Tyb crania (points at right of Fig. 3C, blue colour on the right of Fig. 3I and in Fig. 3J). These maps also showed a smaller snout in Dp1Tyb mice as a result of a reduction in size of the nasal bones (Fig. 3G, H). Although not evident in the heatmaps, the Dp1Tyb cranium was wider as can be seen by the displacement of the zygomatic processes’ landmarks laterally (Fig. 3C and D). The reduced snout and facial widening in combination with the overall smaller size of the Dp1Tyb crania mimics the “mid-face hypoplasia” of human DS. The Dp1Tyb mandibles had a small shape change in the alveolar ramus region and the condylar process but these mandibular changes were all extremely subtle (Fig. 3E, F, K, L).

**Figure 3.**
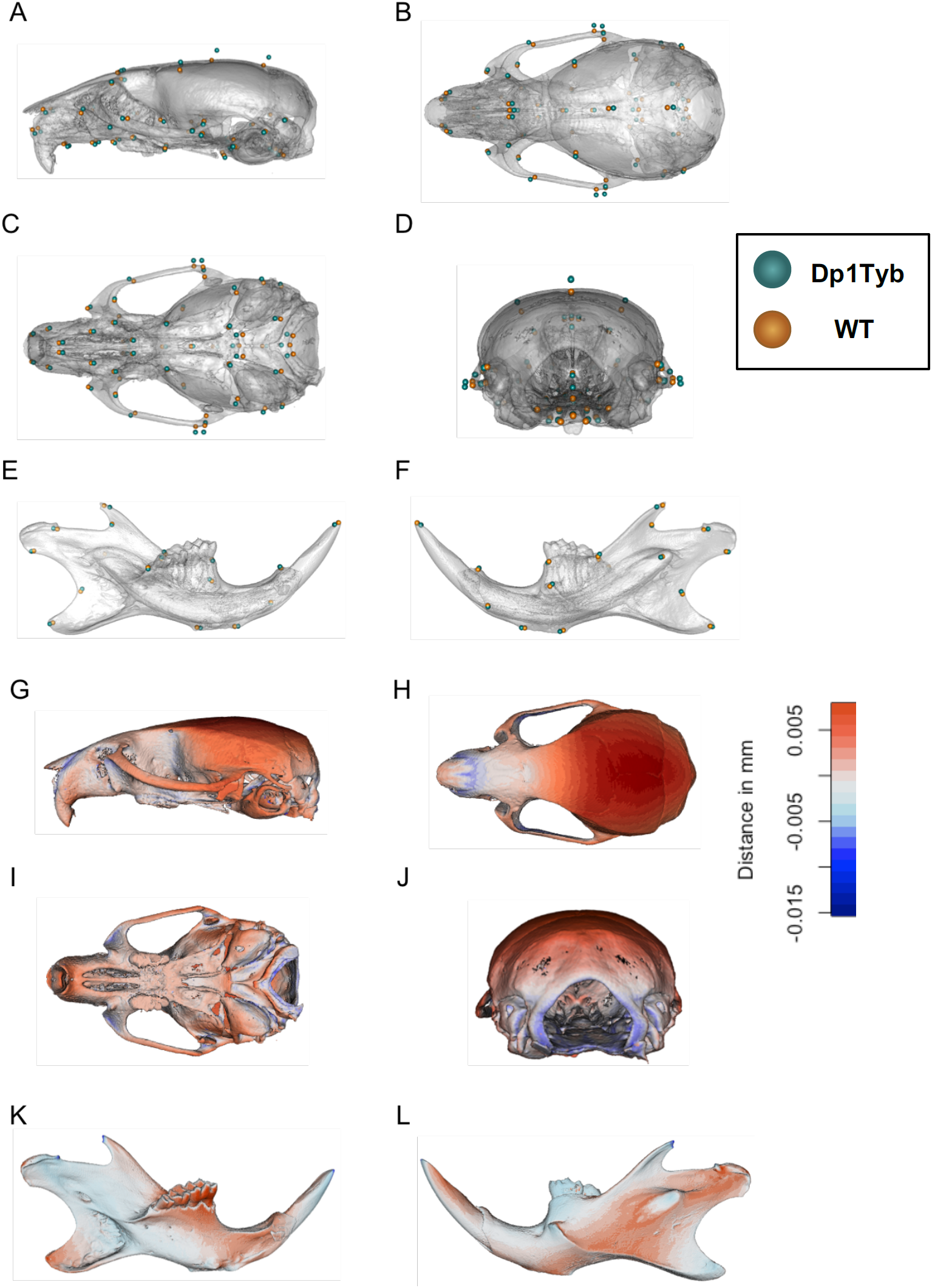
Visualisation of altered shape of Dp1Tyb crania and mandibles determined using landmark-based analysis. **A-F** Mean landmark configurations of WT (Orange) and Dp1Tyb (Cyan) crania (**A-D**) and mandibles (**E, F**), showing lateral (**A**), superior (**B**), inferior and rear (**D**) views of the cranium and lingual (**E**) and buccal (**F**) views of the mandible. **G-L** Displacement heatmaps after global size differences have been regressed out, produced by superimposing the mean shapes of the WT and Dp1Tyb crania (**G-J**) and mandibles (**K**,**L**), showing lateral (**G**), superior (**H**), inferior (**I**) and rear (**J**) views of the cranium and lingual (**K**) and buccal (**L**) views of the mandible. Red and Blue represent the distribution of expansion and contraction respectively in Procrustes (shape) distance. Regions of the Dp1Tyb mesh outside the WT mesh are coloured red, whereas any parts inside the WT mesh are coloured blue, thereby showing displacement relative to WT.

Next we made heatmaps based on the higher-resolution landmark-free method. Displacement maps together with the morphing movies visualised the distances between the two mean meshes. We plotted net displacement rather than movement towards or away from the shape centroid so that only one colour was needed in the maps, using videos to show the direction of differences. The landmark-free analysis showed changes mostly similar to those found using the landmark-based method including the same relative doming of the neurocranium, and shortening of the nasal and maxillary processes (Fig. 4A-D, Supplementary Video 3).

**Figure 4.**
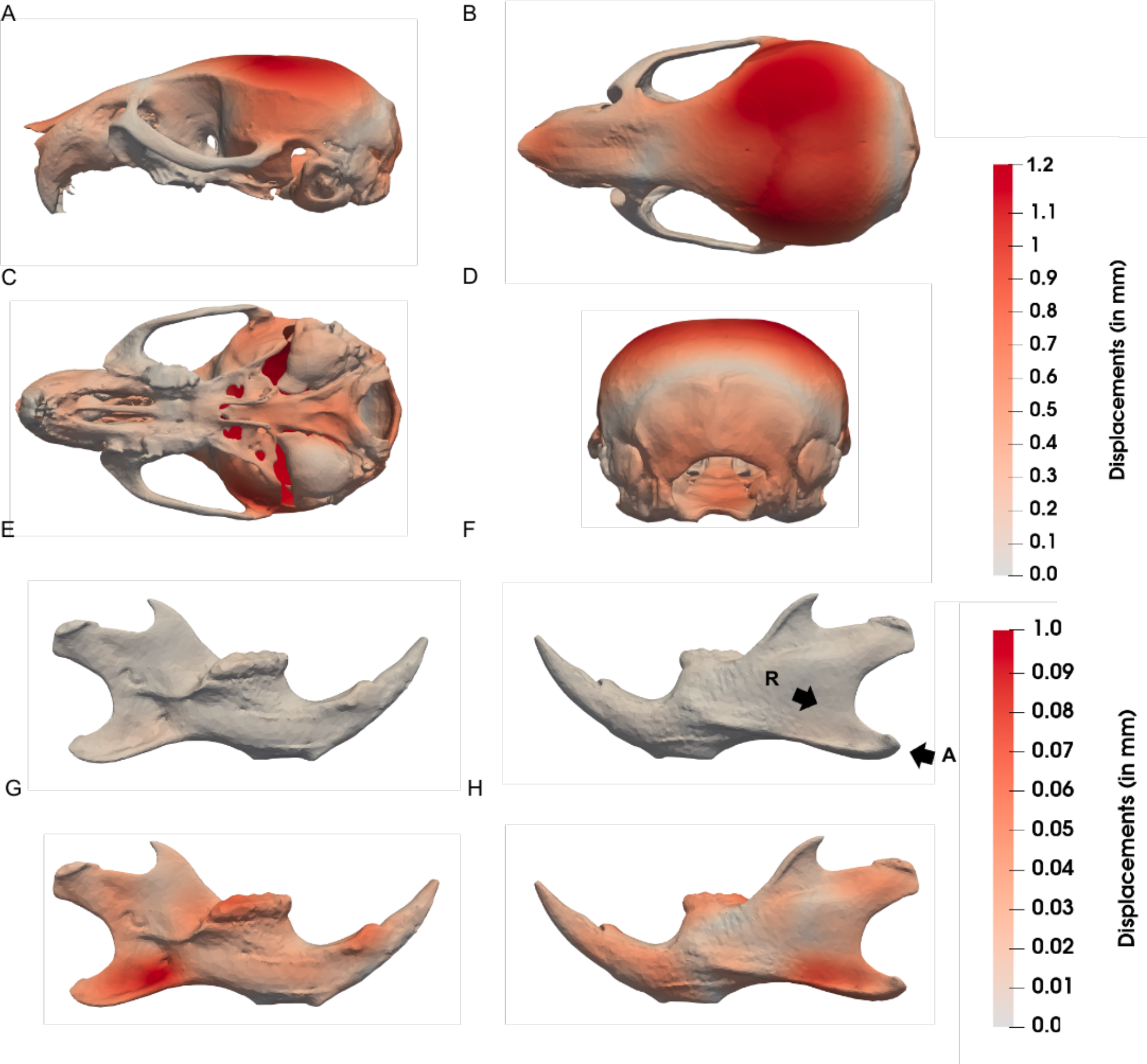
Visualisation of altered shape of Dp1Tyb crania and mandibles determined using landmark-free analysis. Displacement heatmaps after global size differences have been regressed out estimated using output momenta from the current-based atlas construction, showing locations of shape differences between WT and Dp1Tyb crania (**A - D**) and mandibles (**E-H**), showing lateral (**A**), superior (**B**), inferior (**C**) and rear (**D**) views of the cranium and lingual (**E, G**) and buccal (**F, H**) views of the mandible. **A-F** are on the same colour scale; the colour scale in **G, H** maximises contrast for the mandibles only. Arrows **R** and **A** indicate the ramus and Angular process respectively.

Differences between Dp1Tyb and WT mandibles were overall much more subtle (Fig. 4E,F, Supplementary Video 4) consisting of a few tens of microns only. Re-scaling the heatmap colour range better visualised the localisation of the differences (Fig. 4G,H) as did generating a video in which the shape change of the mandible was exaggerated by a factor of three (Supplementary Video 5). Thus, we see the expansion buccally (cheek-wards) of the inferior portion of the ramus (the region just posterior to the molars), the contraction of the angular process, lingual movement of the molar ridge and widening of the incisor alveolus. The expansion of the ramus had not shown up in the landmark-based method because there were no landmarks in this region.

### New shape change information: The landmark-free method maps in-plane deformation

One of the reasons for separating shape difference from size difference in morphometrics is that a simple uniform scale change would appear, artefactually, as change localised distal to whatever point was used as the common frame of reference (i.e. if the centroids are used, there is an increasing centre-to-edge gradient – see demonstration in Suppl. Fig.3). Scaling avoids this problem, but throws away the “ground truth” of the differences. One solution is to find a way of showing size changes entirely locally, capturing surface “stretch” as a measure of local growth differences between specimens. This is also likely to reflect real biological differences which arise in development due to different localised growth. This is not possible with landmarks, but can be done within the high-resolution landmark-free method where a high density of control points is used to guide an even higher density of mesh vertices. Thus, we calculated and mapped local differences in mesh vertex spacing. We used the spacing to generate a heat map without the need for scaling. The results are shown in Fig.5A-F and Supplementary Videos 1, 2 and 6 (see also Supplementary Video 10 for another way of displaying the data). These maps clearly show that the phenotype is almost entirely a growth deficit in the occipital region, posterior of the auditory bulla, facial bones and hard palate. Expansion in the mid-cranial vault is minimal. This representation can be compared with the heatmaps generated on size-scaled data (Fig.5G-L and Supplementary Videos 8 and 9): in these the colour emphasises the expansion of the cranial vault, but this is a relative rather than absolute expansion and does not correspond to actual growth.

**Figure 5.**
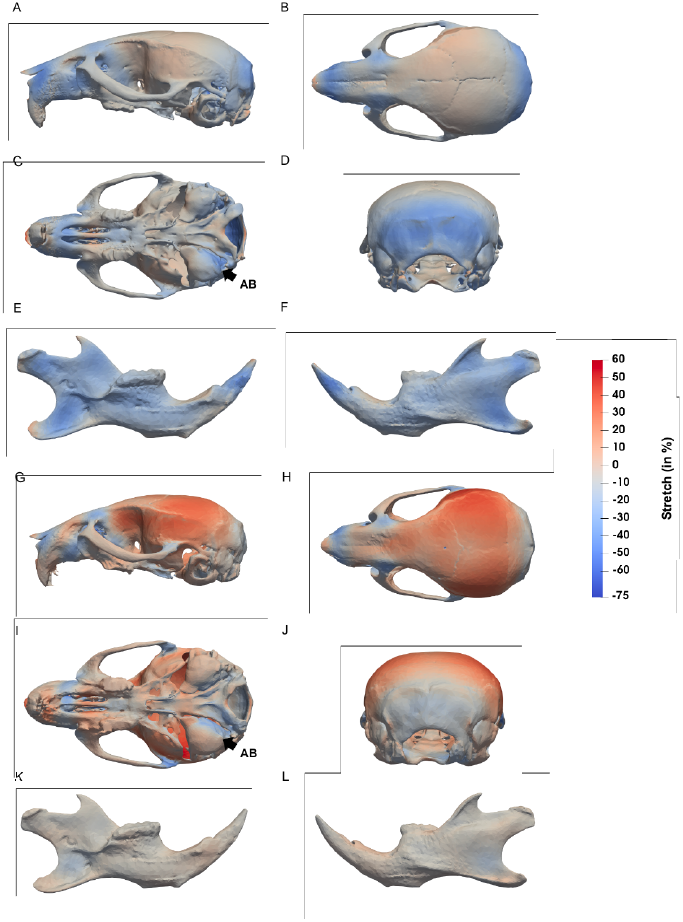
Altered surface stretch in Dp1Tyb crania and mandibles determined using landmark-free analysis. Heatmaps show unscaled (**A-F**) or scaled (**G-L**) surface stretch between WT and DS crania (**A-D, G-J**) and mandibles (**E-F, K-L**), showing lateral (**A**,**G**), superior (**B**,**H**), inferior (**C**,**I**) and rear (**D**,**J**) views of the cranium and lingual (**E**,**K**) and buccal (**F**,**L**) views of the mandible. Stretch changes were estimated using output momenta from the current-based atlas construction. Note that global size changes were regressed out. Arrow **AB** indicates the auditory bulla.

## Discussion

In this paper, we have presented an adaptation and incorporation of a tensor field-based landmark-free shape comparison methodology (Durrleman et al., 2014) into a pipeline that can compare shapes and provide statistical and other data analyses comparable to more traditional geometric morphometrics but with less need for expert training, less labour, less chance of operator error and a higher spatial resolution. We used this pipeline to analyse the previously unexamined craniofacial phenotype of Dp1Tyb mice, a relatively new model of DS (Lana-Elola et al., 2016) and identified localised differences more precisely than was previously possible. The high resolution enabled local deformation density mapping which bypasses some of the issues around global scaling for shape comparison and enabled mapping of the dysmorphology specifically to the occipital and naso-palatal regions.

Both methods revealed that Dp1Tyb mice have size and shape differences compared to WT that parallel the DS phenotype in previously described DS mouse models and in humans. Both separated Dp1Tyb from WT in shape space using one or two principal components and both revealed the significantly reduced size of the cranium and mandible of Dp1Tyb mice, and more specifically, both described brachycephaly (shortened head), resolved in the analysis to an overall size reduction plus cranial doming.

Nonetheless, the landmark-free method has clear advantages. Methodologically, there are three. First is consistency: the landmark-free method overcomes the limitations of manual placement of landmarks which is susceptible to inter- and intra-operator error (Robinson and Terhune, 2017). Second is labour saving: although the landmark-free method requires some manual input in the early stages, particularly in determining image thresholding and in cleaning up imperfect anatomical segmentation, it is substantially less labour-intensive and requires less user training than the landmark-based approach. Third, and perhaps most scientifically significant, is resolution: the landmark-free method provides much higher resolution and information density than the landmark-based method. In principle, the landmark-free method offers arbitrarily high resolution. In practice we found that decimating the initial mesh from ∼2,800,000 to ∼19,000 vertices for the cranium (∼200,000 to ∼16,000 for the mandible) and using a kernel size in the Deformetrica algorithm of 1mm to yield ∼ 2500 control points for the cranium (∼700 for the mandible) captured the interesting anatomical features at high density while avoiding noise, e.g. trivial surface texture differences. Different sizes of specimen will have different optimal spatial parameters. The high density of control points was further refined by having them clustered algorithmically at regions of high variability between samples. This might be contrasted with the inherent bias in landmarking that tends to place shape differences close to landmarks (observable in, for example, Fig. 3A, B). It can also be contrasted with the use of semi-landmarks, which adds landmarks that are evenly distributed across contours or surfaces that are bounded by identifiable landmarks (Adams and Collyer, 2018; Gunz et al., 2005) but still potentially leaves gaps where landmarks are sparse. A trade-off for all methods that increase the number of discrete observations, however, is the increasing as risk of overfitting as the number of variables (e.g. landmarks or voxels) increases relative to the statistical degrees of freedom, or the “curse of dimensionality” (Indyk and Motwani, 1998). Controls for overfitting, such as the permutation test we applied, are therefore essential.

Is the higher resolution and more complete coverage useful? We found that the landmark-free approach allowed us to see changes not visible using the landmark-based approach. Most strikingly, we were able to observe a shape difference in the lower-posterior mandible, where landmarks are absent, and in the snout and palate, where landmarks are more abundant but possibly not dense enough to capture the localised in-plane differences. These latter changes in particular indicate homology with the mid-face hypoplasia found in humans with DS. This will be useful in understanding how DS genes result in dysmorphology because we now have a better knowledge of their location of action.

At another level, high resolution is useful because it enables mapping of surface “stretch” by retaining all the vertices of the mesh, in effect making each vertex a landmark. The very short spatial scale of this mapping is likely to be a much better way to capture and localise changes in biologically causal processes, such as cell proliferation or extracellular matrix expansion, compared to net displacement, where the cause could be hundreds of cell diameters away. The local nature of this deformation mapping makes it easier to interpret the visual display of the deformations without global scaling.

Although the landmark-free method was developed for MRI scans of brains^24^ applying it to a mutant skull has enabled two important conclusions. The first is that using a landmark-free approach is still advantageous even when traditional landmarking is possible. The second conclusion is that it is possible to apply this approach in the form of a relatively user-friendly tool. We have found it useful in understanding the DS craniofacial phenotype but, with modest computational expertise, other researchers can tackle any mutant phenotype, including where traditional methods have struggled, such as in early developmental stages or other biological forms that lack well defined landmarks.

## Materials and Methods

### Mice and imaging

C57BL/6J.129P2-Dp(16Lipi-Zbtb21)1TybEmcf (Dp1Tyb) mice (Lana-Elola et al., 2016) were bred at the MRC Harwell Institute. All mice were backcrossed to C57BL/6JNimr for at least 10 generations. All animal work was approved by the Ethical Review panel of the Francis Crick Institute and was carried out under Project Licences granted by the UK Home Office. Heads from 20 mice at 16 weeks of age were used (10 Dp1Tyb and 10 WT, 5 male and 5 female of each genotype). However, 4 subjects were excluded from the analysis due to fractures in either the mandible or skull. Heads were prepared for micro-computed tomography (µCT) by fixation in PFA and then scanned at a 25 µm resolution using a µCT 50 (Scanco).

### Landmark-based morphometric analysis

#### Landmark Acquisition

Three-dimensional locations of 68 anatomical landmarks for the cranium and 17 landmarks for the mandible (Supplementary Fig. 1) as previously defined by Hallgrimsson and colleagues (Hallgrimsson et al., 2007) were placed onto 3D reconstructions of µCT images using Microview (Parallax Innovations). Landmarks were placed manually and verified by checking orthogonal planar views of the subject.

#### Statistical analysis of landmark dataset

Landmarks for all subjects were aligned using Procrustes superimposition, and distances between landmarks for each subject were analysed by Principal Component Analysis (PCA) using MorphoJ (Klingenberg, 2011) to visualise group separation by shape. To quantify significance in shape differences between genotypes we used the Procrustes Distance Multiple Permutations test (1000 iterations) within MorphoJ. Centroid size was calculated as the square root of the sum of the squared distances from each landmark to the centroid i.e. the centre of mass of all landmarks of a given specimen and was used to compare size differences between WT and Dp1Tyb crania and mandibles. The statistical significance of such size differences was calculated using a 2-tailed unpaired t-test.

Shape changes after Procrustes scaling were visualised as heatmaps. Using the Morpho R package function tps3d (documentation for which can be found at https://www.rdocumentation.org/packages/Morpho/versions/2.6/topics/tps3d), mean landmark sets of each genotype group were used to interpolate an average mesh, using a thin-plate spline method (Bookstein, 1989), thus providing a mesh for both the mean WT shape and mean Dp1Tyb shape. The Morpho R meshdist function (https://www.rdocumentation.org/packages/Morpho/versions/2.6/topics/meshDist) was then used to create the heatmaps. The function first calculates the distances of the reference meshes vertices (in this case mean WT mesh) to that of the target mesh (mean Dp1Tyb mesh). Then, using a previously proposed algorithm (Baerentzen, 2005), the distances were given a negative value if inside the reference mesh or a positive value if outside. A vector containing blue and red colour values was assigned to the negative and positive values respectively (Schlager, 2017).

### Landmark-free morphometric analysis

As an alternative to landmark population comparisons, statistical analysis of anatomical shapes can be achieved using so-called atlas-based approaches, which consist of estimating an anatomical model (i.e. template) as the mean of a set of input shapes (rather than point clouds) and quantifying its variation in a test population as deformations. This was previously achieved in a reproducible and robust landmark-free manner by Durrleman et al (2014). This approach bypasses a number of problems associated with mesh point-to-point comparison by representing deformation between shapes as the diffeomorphic transformation of flow fields, i.e. currents over the mesh surface. Currents are parameterised by a set of control points in space and initial velocities, or momenta. By means of a gradient descent optimisation scheme, the method is able to produce a statistical atlas of the population of shapes. An atlas refers to a mean template shape, a set of final control point positions, and momenta parameterising the displacements between the template to each initial individual shape. In the following sections we describe the different steps to achieve such analysis.

#### Landmark-free Atlas Construction

Step 1 (Fig. 1, Table 1, Supplementary Appendix 1). Despite the fact that images acquired using µCT show good bone contrast, they often include the presence of artificial objects (noise and debris in the specimen), small holes and cartilage that need be excluded in order to obtain consistently comparable final surface meshes. To extract the surface meshes, a series of image processing steps were applied. After a thresholding operation to extract the skull, “morphological opening” and “closing” were performed on the binary mask to remove internal cartilage structures (Supplementary Fig. 2). Removal of spurious objects was achieved by clustering, categorising all of the connected components in an image by size and retaining only the largest component. Skulls were segmented using bone density to isolate the mandible (whose density is higher than that in the rest of the skull). However, this segmentation of the mandible can happen improperly and may include parts of the temporal bone which must be cleaned and removed manually. Mandible meshes were extracted from the skull binary mask semi-automatically using Watershed segmentation (Mangan and Whitaker, 1999).

Step 2. The meshes were produced using marching cubes on the binary images, followed by a surface Laplacian smoothing (Vollmer et al., 1999) and finally decimation of a factor of 1/80 to decrease mesh resolution and reduce overall computation time.

Step 3. The atlas construction necessitates the production of aligned meshes from the input µCT images as a pre-processing step. Minimally 3 pairs of landmarks were chosen to align meshes using a Procrustes technique (Gower, 1975) using a rigid body-plus-scaling transformation model (similarity alignment) or a rigid body transformation without scaling (rigid body alignment).

Step 4. As previously described (Durrleman et al., 2014), the atlas construction major hyper-parameters consist of (1) the size of the Gaussian kernel used to represent shapes in the varifold of currents, denoted σ_W_, and (2) the number of control points, denoted N_cp_. Control points can be thought of as unbiased landmarks, initially they are spaced on a regular grid evenly but move to areas of greatest variation. Thus as much as possible of the shape change is captured in an unbiased manner. σ_W_ can be seen as the precision at which the shape deformation is described. N_cp_ denotes the sampling density in space. In our experiments, we fixed σ_W_ = 2 mm. Using that scale, the derived number of control points Ncp was 350 and 940 for the mandibles and the cranium analyses respectively. All subjects’ meshes were used for the construction of the atlas. The optimisation procedure typically converged within 150 iterations in approximately 20h on a 10 core cpu for the mandible. The atlas construction produces the following outputs:

- **Control Points**: For each subject, the N_cp_ final control points locations {c}, that correspond to areas that describe the population’s most important variability in shape.
- **Momenta**: For each subject, the N_cp_ final momenta {m} associated to each control point, that describe the shape directional variation from the population average.
- **Template**: Mean shape of the population.

#### Landmark-free dataset statistical analysis

Step 5 (Fig. 1). The atlas outputs provide a dense amount of information that can be used for various statistical analyses. Centroid size was calculated as the square root of the sum of the squared distances from each mesh node to the centroid of all nodes of a given specimen. Non-linear Kernel PCA with dimension 5 was applied to the set of momenta produced from the atlas of the population, in order to find the principal modes of variation of the entire population. The resulting output provided a way to compare these results with the landmark-based PCA analysis. We projected the subjects onto the feature space for comparison purposes. Such projection provides dense information of shape differences between the two sub-populations. Local magnitude of the momenta interpolated at the template mean mandible (or cranium) mesh point locations allow for additional qualitative interpretation of shape differences between groups. Stratified k-fold cross-validation analysis was performed on the PCA data to evaluate the statistical power of classification between the two groups. Significance of the classification score was tested using a multiple permutations test at 1000 iterations (Ojala and Garriga, 2010). To assess overfitting, subjects were randomly partitioned into two groups (“scrambled groups”) and PCA analysis performed to generate inter-group vectors. The distribution of the vector magnitudes was tested for Normality using the Shapiro-Wilk test.

## Code availability

Python scripts and documentation for the landmark-free morphometric analysis are freely available on GitHub (https://gitlab.com/ntoussaint/landmark-free-morphometry) and their use is also described in Supplementary Appendix 1.

## Acknowledgements

We thank Heather Cater and Sara Wells from MRC Harwell for breeding the mice and Chris Healy at KCL for performing the micro-CT scans.

## Author contributions

NT designed and wrote the code for the analysis. YR and WL carried out experiments. NT, YR and WL analysed the data. YR, NT, VLJT, JAS and JBAG wrote paper. BH, EMCF, VLJT, JBAG and JAS supervised the work.

## Supplementary Information

### Supplementary Videos

**Supplementary Video 1.**

**Cranium morph including size**

Cranium morph between the mean WT and Dp1Tyb shapes, where size has not been regressed out. Derived from landmark-free analysis. Red and Blue colour change indicate expansion and contraction respectively as a percentage. Derived from landmark-free analysis.

**Supplementary Video 2.**

**Mandible morph including size**

Mandible morph between the mean WT and Dp1Tyb shapes, where size has not been regressed out. Red and Blue colour change indicate expansion and contraction respectively as a percentage. Derived from landmark-free analysis.

**Supplementary Video 3.**

**Cranial morph displacement heatmap**

Cranial morph between the mean WT and Dp1Tyb shapes, where size has been regressed out. Red colour change indicates the magnitude of displacement in mm. Derived from landmark-free analysis.

**Supplementary Video 4.**

**Mandible morph displacement heatmap**

Mandible morph between the mean WT and Dp1Tyb shapes, where size has been regressed out. Red colour change indicates the magnitude of displacement in mm. Derived from landmark-free analysis.

**Supplementary Video 5.**

**Exaggerated Mandible Morph**

Mandible morph between the mean WT and Dp1Tyb shapes, where size has been regressed out and the magnitude of the deformation has been exaggerated by a factor of 3. Derived from landmark-free analysis.

**Supplementary Video 6.**

**Cranial Interior morph including size stretch heatmap**

Cranial morph, showing the interior of the skull, between the mean WT and Dp1Tyb shapes, where size has not been regressed out. Red and Blue colour change indicate expansion and contraction respectively as a percentage. Derived from landmark-free analysis.

**Supplementary Video 7.**

**Cranial Interior morph stretch heatmap**

Cranial morph, showing the interior of the skull, between the mean WT and Dp1Tyb shapes, where size has been regressed out. Red and Blue colour change indicate expansion and contraction respectively as a percentage. Derived from landmark-free analysis.

**Supplementary Video 8.**

**Mandible morph stretch heatmap**

Mandible morph between the mean WT and Dp1Tyb shapes, where size has been regressed out. Red and Blue colour change indicate expansion and contraction respectively as a percentage. Derived from landmark-free analysis.

**Supplementary Video 9.**

**Cranial morph stretch heatmap**

Cranial morph between the mean WT and Dp1Tyb shapes, where size has been regressed out. Red and Blue colour change indicate expansion and contraction respectively as a percentage. Derived from landmark-free analysis.

**Supplementary Video 10.**

**Cranial mesh points stretch heatmap**

Landmark-free-derived heatmap of cranium local stretch with no scaling. The mesh is represented here as all the points of the mesh rather than a rendered surface. Red and Blue colour change indicate expansion and contraction respectively. Downloading and viewing in a paused video by moving the slider is recommended.

### Supplementary Figures

**Supplementary Figure 1.**
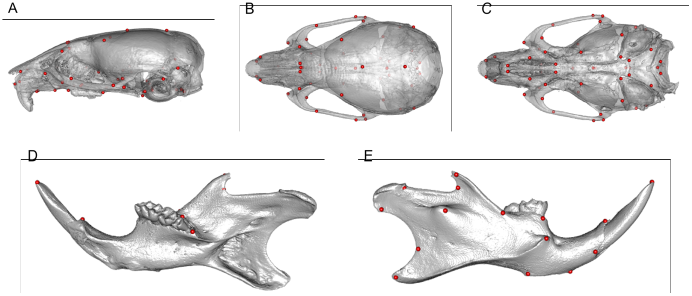
Location of landmarks used in landmark-based morphometric analysis. **A-E** Anatomic locations of landmarks shown on a 3D digitized image of a µCT scan as previously defined ^1,2^. 68 cranial landmarks are shown on lateral (A), superior (B), and inferior (C) views of the cranium. 17 mandibular landmarks are shown on lingual (D) or buccal (E) views of the mandible.

**Supplementary Figure 2.**
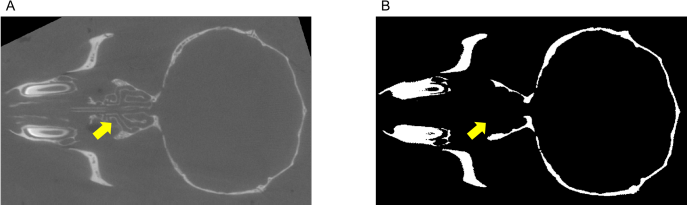
Planar view of mouse head captured by µCT. **A** Image pre-processing. **B** Image post-processing. Yellow arrow indicates cartilaginous element (nasal turbinate).

**Supplementary Figure 3.**
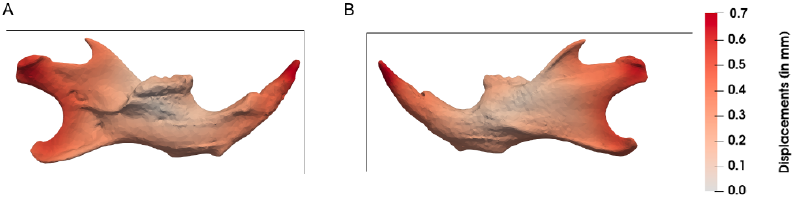
Displacement heat map with no scaling. Without scaling, displacement changes are localised to the extremities.

### Supplementary Table

**Supplementary Table.** Comparison of centroid sizes of crania and mandibles from WT and Dp1Tyb mice determined using landmark-based or landmark-free methods.

| | | Mean Centroid Size in mm<br>(± SEM) | | difference | |
| --- | --- | --- | --- | --- | --- |
|  |  | WT | Dp1Tyb | absolute (mm) | percentage |
| Landmark-based | Cranium | 6.387±0.061 | 5.939±0.061 | 0.448 | 7.01% |
| | Mandible | 4.502±0.038 | 4.204±0.038 | 0.298 | 6.62% |
| Landmark-free | Cranium | 6.987±0.063 | 6.514±0.063 | 0.473 | 6.77% |
| | Mandible | 4.025±0.041 | 3.768±0.041 | 0.257 | 6.39% |

## Supplementary Appendix 1

### Landmark-free Morphometrics Pipeline Description

This Appendix is intended as an overview of the landmark-free pipeline for a relative non-specialist. Detailed dcoumentation is embedded with the pipeline code published on GitLab at https://gitlab.com/ntoussaint/landmark-free-morphometry. This Appendix is organised into five sections as follows:

A. Pipeline Dependencies
B. µCT Pre-processing
C. Mesh Alignment
D. Atlas Construction
E. Shape Statistics

#### A. Pipeline dependencies and notes on initialisation

All packages and software listed below are required. The pipeline is written in Python and C++ and requires a suitable environment to run both. We used the browser-based Juypiter environment (https://jupyter.org/). Specific requirements for each segment of the pipeline are listed at the appropriate page within the pipeline code but the complete list is as follows:

Absolutely required:

**FSL -** https://fsl.fmrib.ox.ac.uk/fsl/fslwiki (Image analysis tools)

**Deformetrica -** http://www.deformetrica.org/ (Atlasing tools)

**Git -** https://git-scm.com/(Code version-control system)

**ITK -** https://itk.org/ (Segmentation and registration tools)

**VTK -** https://vtk.org/ (Visualisation tools)

**CMake -** https://cmake.org/ (Compiler)

**Python packages:**

Matplotlib

Numpy

SimpleITK

Vtk

Pandas

Seaborn

Software we used but for which there may be similar alternatives:

**Meshlab -** http://www.meshlab.net/ (Mesh decimation)

**Itksnap -** http://www.itksnap.org/pmwiki/pmwiki.php (Image manipulation tool)

**Paraview -** https://www.paraview.org/ (Image visualisation tools)

**Mitk -** http://mitk.org/wiki/The_Medical_Imaging_Interaction_Toolkit_(MITK) (Landmarking tool for coarse alignment)

The pipeline code is downloaded as landmark-free-morphometry-master.zip or ∼.tar. After decompressing the package and moving to its directory within the Terminal app (Apple, Linux or equivalent in Windows) Juypiter is started using the command **ipython notebook** which opens the pipeline in the default system Web browser. The pipeline is presented as a series of virtual “pages” which contain “**cells**” (sections) of two types. One type defines a function, the other applies those functions to the data. Cells can be run individually or as an automatic sequence (see Juypiter documentation for details). An example of a function-defining cell is below:

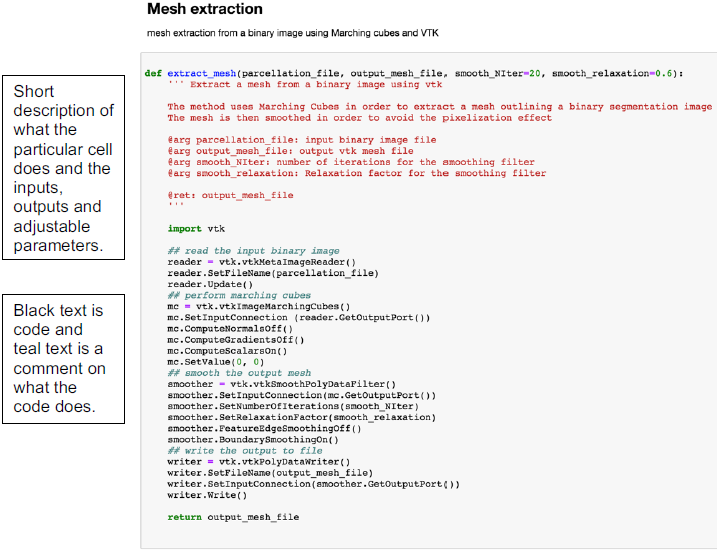

#### B. µCT Pre-processing

This section describes the code used to extract and produce a mesh from an initial µCT image. The following cells can be run with default values to define the required functions:

##### Binary Segmentation

- Defines functions used in the object extraction

##### Parcellation

- Defines functions required for parcellation of binary image

##### Mesh Extraction

- Defines function used in generating a mesh from a binary image

##### Mirror mesh file

- Defines function used to mirror the mesh file

##### Display help functions

- Defines functions used to display images within the notebook

##### Generic Imports

- Imports Numpy

Using **ITK-Snap** save µCT images in the **NIFTI** (.nii.gz) format.

The following five **cells** need to be edited to change parameters and filenames before running (N.B. annotations describe the sections that can or need to be changed, along with a brief description of inputs, outputs and function/s of the cell).

##### Load data

Loads in the image file to be processed.

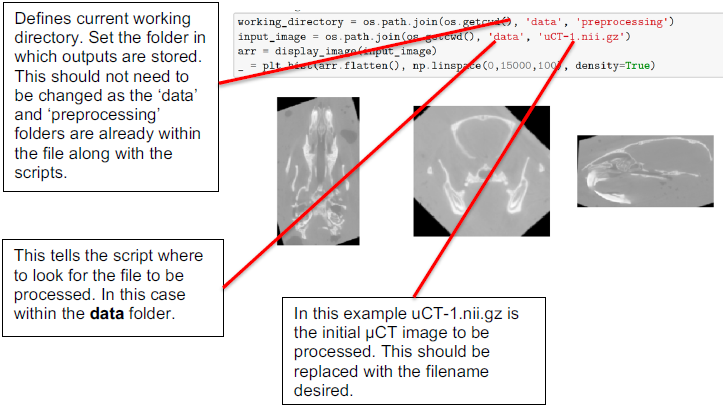

##### Object extraction

The cell uses the functions defined in **object_extraction** to extract the object of interest from the input image and apply morphological functions to remove any noise and close unwanted holes.

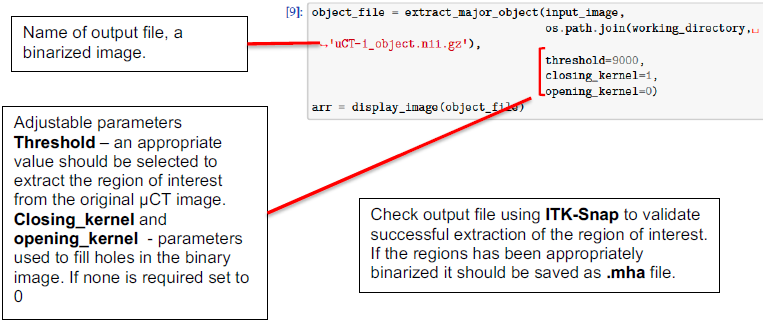

##### Parcellation

This cell uses the functions defined in **parcellation** to ‘parcellate’ the image into various regions according to the density of the bone. This allows the separation of mandible (densest bone) from the rest of the skull. If no region separation is required skip this step and move to the **Mesh extraction** step:

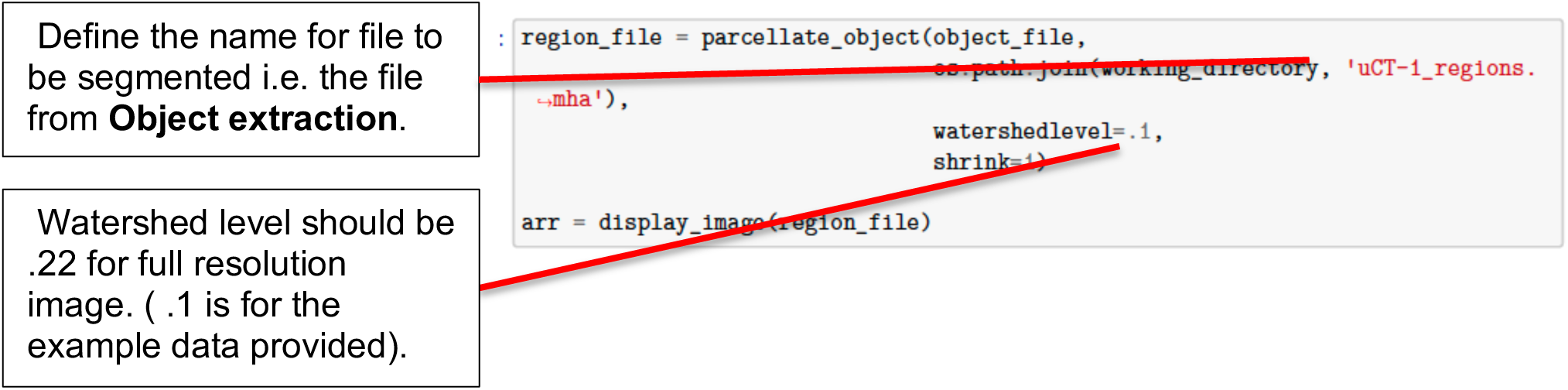

The output is a binary image with regions ordered by bone density. Use **ITK-Snap** to visualise these labels. The example on the right shows these regions, here label one is the red region. Note the label/s for region of interest (ROI).

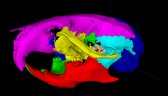

##### Object selection

Input the labels of relevance to separate the ROI from the rest of the subject.

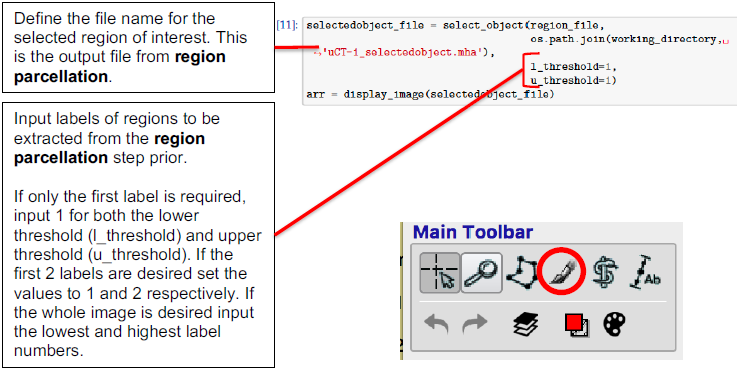

Check the output in **ITK-snap** opening it as a **segmentation file.** Any noise or irrelevant regions can be removed using the paint brush tool.

##### Extract mesh

This step uses functions defined in **Mesh Extraction** to extract a surface mesh from the region of interest selected prior. The meshes generated are in the **.vtk** format.

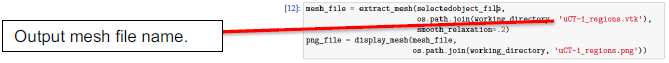

Mesh files produced are large in size and increase computational time of the **Atlasing** step to unrealistic durations. To overcome this, meshes were decimated using **meshlab**.

Save meshes in the **.stl** (stereolithography) format using paraview. Open the mesh in meshlab and apply **quadric edge collapse decimation** fi and enter the percentage reduction desired (authors used 0.15).

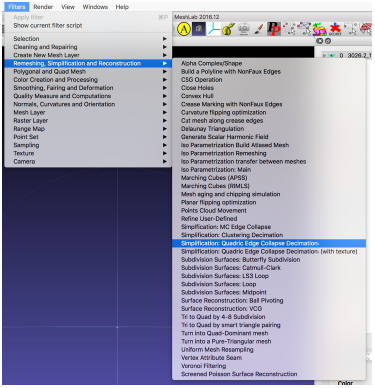

Once meshes for all subjects are generated move to the next step, **Mesh Alignment.**

#### C. Mesh Alignment

This section describes the process of aligning the meshes produced in the previous section. Initial coarse alignment is required to allow the Atlasing process (below) to work. This first requires manually placing a small number of landmark using **Mitk-Workbench**, minimally three (e.g. for a hemi-mandible) or more usually three symmetrically placed pairs. Landmark file names should be identical to the mesh file name e.g. Mandible_1.vtk, with its landmark file, Mandible_1.mps.

Below is shows landmarks used to align the mandible.

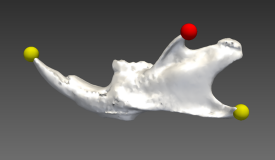

Once complete move landmark files (.mps) and mesh files (.vtk) into Alignment folder. The following are run with default values:

##### Read landmarks

- Defines functions used to import landmark files

##### Numpy – vtk help functions

- Define functions used convert landmark files so they can be used to align meshes
- Functions used to extract centroid and centroid size

##### Procrustes alignment

- Defines functions used in alignment
  - Rigid – rigid body alignment
  - Similarity – rigid body plus scaling alignment

##### Load data

Loads meshes and associated landmark files ready for alignment,

##### Mesh Alignment

Uses functions defined in **Procrustes alignment** to align meshes. Two modes can be selected, **Similarity** or **Rigid body**. “Similarity” aligns meshes whilst regressing out the size. “Rigid body” aligns meshes but maintains size differences:

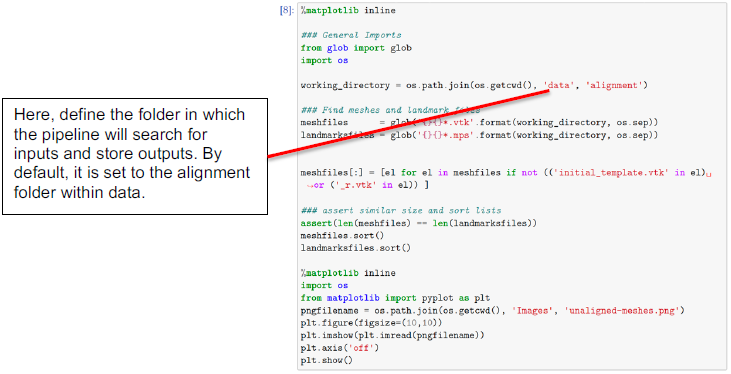

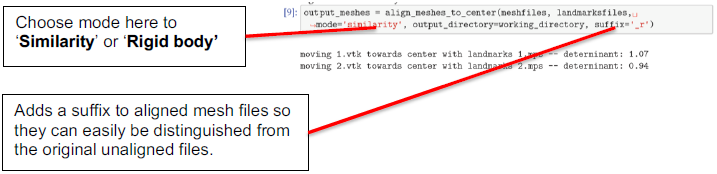

Mesh alignment fidelity should be checked in **Paraview**.

#### D. Atlas Construction

This section describes the application of the shape averaging and comparison (atlasing) processes.

Within the **atlasing** folder (included in the pipeline package) there are three files adjustable files:

**Model.xml** describes parameters for deformation:

- kernel-width: Order of magnitude of displacements for the template
- kernel-width: Order of magnitude for the subjects’ displacements

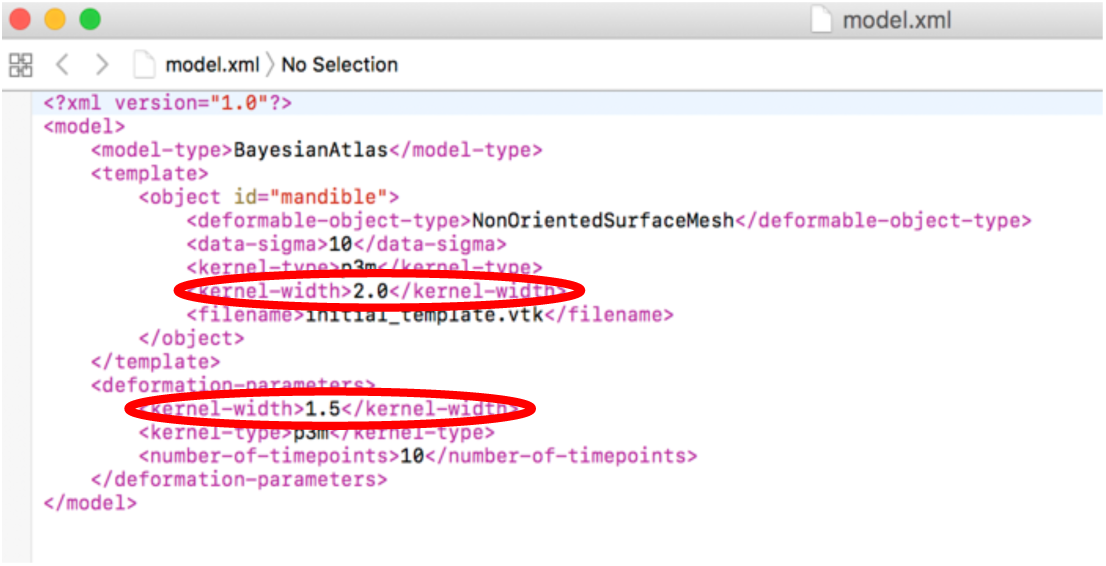

**Optimization_parameters.xml** describes parameters that drive the minimization procedure:

- max-iterations: Stop criterion for iterations, usually ∼150
- number-of-threads: Number of threads to use, usually equal to the number of subjects (should not exceed the number of threads available on the system)

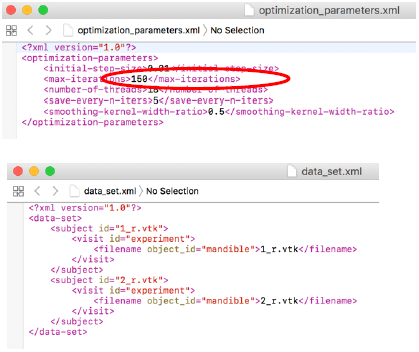

**Data_set.xml** defines the files to be atlased and should be structured as follows:

From the alignment folder transfer aligned meshes (files with suffix **_r**) and initialtemplate.vtk into the atlas folder.

##### Deformetrica

The first cell loads the **Deformetrica** software. If the software is not found an error is returned.

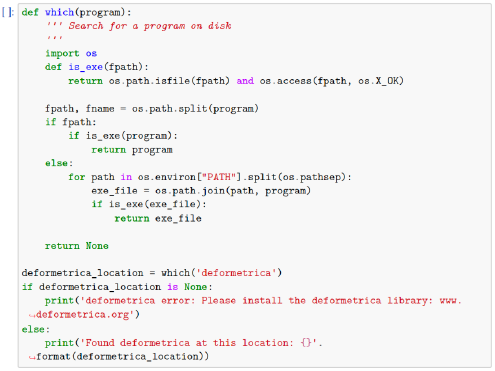

##### Launch Simulation

Launches the atlasing software using the files and parameters defined by model.xml, optimisation_parameters.xml and data_set.xml.

**Deformetrica.log**, within the output folder, contains information on the progress of the atlas, including the number of control points and the current iteration.

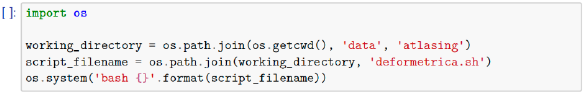

#### E. Shape statistics

This section describes the statistical analysis of the outputs from the atlas.

Atlasing generates several output files of which the following three are used for statistical analysis:

- **Atlas_controlpoints.txt**
  - Control points of the atlas

- **Atlas_momenta.txt**
  - Momentum vectors of the atlas

- **Atlas_initial_template.txt**
  - Average mesh of the population

These files are must be moved into the **shape statistics** folder. In the same folder, a user-populated file **data.csv** defines the names and types of the subjects.

**data.csv** example file

In this example in GroupId, 1 refers to a mutant and −1 to WT, and in Gender, 1 Male and −1 female.

|  | A | B | C |
| --- | --- | --- | --- |
| 1 | id | GroupId | Gender |
| 2 | 33.4a | -1 | -1 |
| 3 | 33.4b | -1 | -1 |
| 4 | 35.1c | -1 | -1 |
| 5 | 37.1c | 1 | 1 |
| 6 | 38.1c | -1 | 1 |
| 7 | 29.3d | -1 | 1 |
| 8 | 29.3g | -1 | 1 |
| 9 | 30.3b | -1 | -1 |
| 10 | 30.3d | 1 | -1 |

The following cells are the run with default values:

##### Imports

- Loads relevant packages required for this section of the pipeline

##### Deformetrica

- Loads deformetrica ready to be used (required for creation of morphs)

##### Load data

- Loads output files stated previously to be analysed

##### Define population’s groups

- Loads data.csv

##### Subgroup definitions

- Assigns names to groups of specimens defined by one or more parameters in **data.csv**.

[Default groups are: WTf (Wildtype female, defined by: group ID = −1, gender = −1, CRf is defined by: group id = 1, gender =-1), WTm (Wildtype Male), CRf (Carrier female) and CRm (Carrier Male)]

However, the user should change the names and definitions of subgroups to suit their needs.

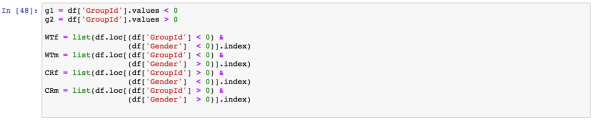

##### Kernel Principal Component Anaylsis (kPCA)

This cell runs a principal component analysis (PCA) and produces a classification score (as a percent) and a p value for said score. As well as defining group IDs (by default group one > 0 and group two < 0 using values set in **data.csv**) and calculating the mean Principal Component score.

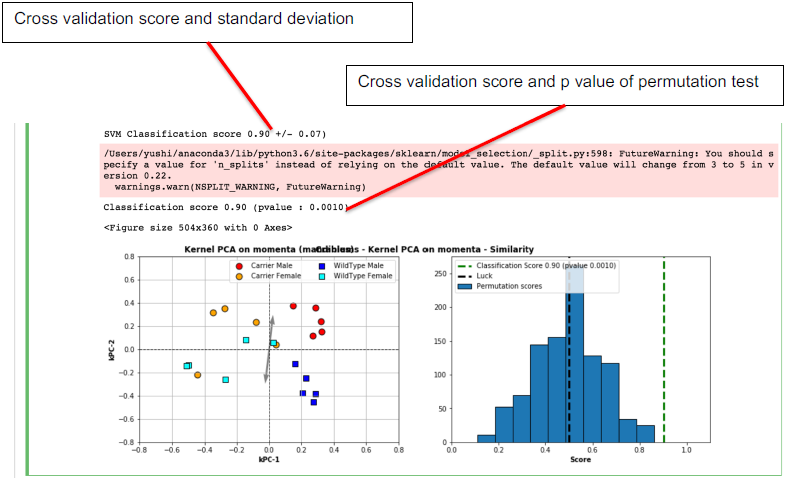

##### Eigenvalues/Eigenvectors

Produces Eigenvalues and eigen vectors. Providing data on how much variability each principle component describes.

##### Write PCA points on disk

Produces an excel file, kpca.csv containing the PC values which can exported to produce PCA graphs.

##### Momenta projection

Produces momenta that describes project between the mean of the whole population to the means of the previously defined populations.

##### Shoot mean shape between groups

Uses the momenta projection to ‘shoot’ the shape towards the means of the two groups.

##### Shooting outputs

Saves the shot meshes produced in the previous step in two locations:

- data/shapestatistics/shooting/forward
- data/shapestatistics/shooting/backward

They correspond to the mean shape (**Atlas_initial_template.vtk**) displaced towards the mean of group one (backward) and the mean of group two (forward). Each file now contains several meshes. The first mesh in the folder is the average shape of the whole population and the last mesh the mean shape of the subgroup. Each mesh in-between represents a step of deformation moving from one to the other.

##### Cyclic deformations between subgroups

Reorders the mesh files produced in **Shooting outputs** to generate a cyclic shape change starting with the mean of group one to the mean of group two and back again. The files for the cyclic deformation are saved in: data/shapestatistics/shooting/combined. Files should be opened in **Paraview** to visualise the meshes.

The deformation is viewed cyclically, by pressing the repeat and play buttons.

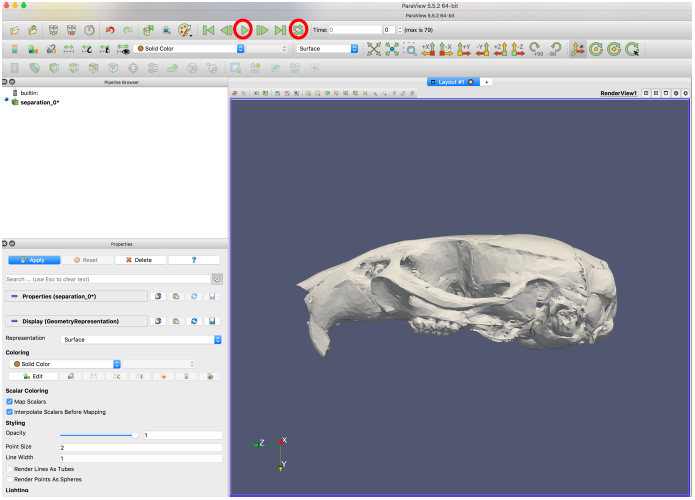

##### “Stretch” map (Local volume change map) between groups

This step integrates data on displacement and stretch into the cyclic deformation produced in the previous step. Removing the # from in front of **compute_point_displacements** calculates displacement.

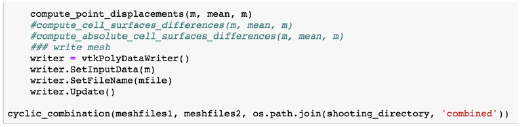

Removing the # from in front of **compute_cell_surfaces_difference**, calculates stretch.

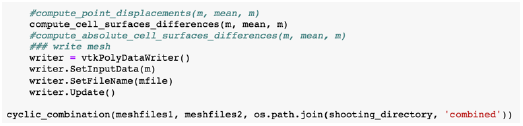

Heat maps are viewed by importing the mesh files from the combined folder into **Paraview**. Select displacements or Absolute Volume from the colouring drop down menu and the morph was played as before to visualise the heat maps. Colour scale (look-up table) may need to be adjusted to visualise differences properly.

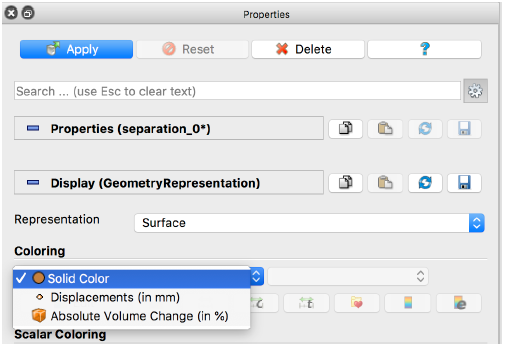

